# A prophage-encoded sRNA limits lytic phage infection of adherent-invasive *E. coli*

**DOI:** 10.1101/2025.05.06.652453

**Authors:** Robert S. Brzozowski, Amelia K. Schmidt, Nicole L. Pershing, Annika Dankwardt, Dominick R. Faith, Alex C. Joyce, Andrew Maciver, William S. Henriques, Shelby E. Andersen, Blake Wiedenheft, Sherwood R. Casjens, Breck A. Duerkop, June L. Round, Patrick R. Secor

## Abstract

Prophages are prevalent features of bacterial genomes that can reduce susceptibility to lytic phage infection, yet the mechanisms involved are often elusive. Here, we identify a small RNA (*svsR*) encoded by the lambdoid prophage NC-SV in adherent-invasive *Escherichia coli* (AIEC) strain NC101 that confers resistance to lytic coliphages. Comparative genomic analyses revealed that NC-SV–like prophages and *svsR* homologs are conserved across diverse Enterobacteriaceae. Transcriptional analyses reveal that *svsR* represses maltodextrin transport genes, including *lamB*, which encodes the outer membrane maltoporin LamB—a known receptor for multiple phages. Nutrient supplementation experiments show that maltodextrin enhances phage adsorption, while glucose suppresses it, consistent with established effects of these sugars on *lamB* expression. *In vivo,* we compared wild-type NC101 and a prophage-deletion strain (NC101^ΔNC-SV^) in mice to assess the impact of NC-SV on lytic phage susceptibility. Although intestinal *E. coli* densities remained stable across groups, animals colonized with NC101 exhibited markedly reduced phage burdens in both the intestinal lumen and mucosa compared to mice colonized with NC101^ΔNC-SV^. This reduced phage pressure was associated with increased dissemination of NC101 to extraintestinal tissues, including the spleen and liver. Together, these findings highlight a nutrient-responsive, prophage-encoded mechanism that protects AIEC from phage predation and may promote bacterial persistence and dissemination in the inflamed gut.

## Introduction

Bacteriophages (phages) are abundant viruses that infect bacteria and play a key role in shaping microbial communities. Their impact is especially significant in densely populated environments like the gut. Many phages exist as prophages—viral genomes integrated into bacterial chromosomes—that often provide competitive advantages to their hosts. One well-established function of prophages is protection against competing phages, enhancing host survival in phage-rich niches [1–9]. However, the specific mechanisms by which individual prophages confer this resistance are highly diverse and not yet fully understood.

Environmental factors such as diet shape the gut microbiota and contribute to dysbiosis in inflammatory bowel diseases (IBD) like Crohn’s disease and ulcerative colitis [10–12]. One example is maltodextrin, a common dietary additive which degrades mucus integrity, exacerbates inflammation [13, 14], and promotes the expansion of adherent-invasive *Escherichia coli* (AIEC) [15, 16], a pathobiont enriched in the IBD gut [10, 11, 17, 18]. Maltodextrin uptake in *E. coli* is mediated by the outer membrane maltoporin LamB, a known receptor for several coliphages, including lambda and others [19–22]. Thus, metabolic responses to dietary components may directly influence phage susceptibility.

As enteric bacteria proliferate in the inflamed gut, so too do phages that target them [6, 7, 23–25]. This expansion of enteric phages can reshape microbial communities and potentially influence IBD progression [24, 26, 27]. Despite this, how pathobionts like AIEC persist and replicate amid rising phage pressure remains unclear.

In this study, we identify a lambda-like prophage, NC-SV, in AIEC strain NC101 that confers protection against some lytic phages by reducing virion adsorption to the bacterial surface. Using RNA sequencing, we found that NC-SV represses maltodextrin transport genes, including the phage receptor *lamB* [21, 22]. Previous studies have shown that maltodextrin upregulates *lamB* expression while glucose represses it via catabolite repression [28]. Consistent with this, we observed that maltodextrin enhances phage adsorption, whereas glucose reduces it. We further identified a prophage-encoded small RNA (sRNA) that downregulates *lamB* and reduces phage adsorption.

Together, our findings reveal a mechanism by which prophages modulate host nutrient uptake and surface receptor expression to protect against viral infection. This strategy may be particularly advantageous to AIEC in the altered nutritional landscape of the inflamed IBD gut [29–33].

## Results

### The NC-SV prophage protects *E. coli* NC101 from lytic phage infection by reducing virion adsorption

Many phages that inhabit the gut reside as prophages integrated into the genome of their bacterial host [34]. The genome of AIEC strain NC101 contains three intact prophages with three distinct morphotypes including Siphovirus (NC-SV), Myovirus (NC-MV), and Inovirus (NC-Ino) (**Table 1**). NC-SV is related to phage lambda and is integrated at the phage 21 attachment site within the isocitrate dehydrogenase gene (*icd*) in the *E. coli* NC101 chromosome. NC-SV encodes lambda-homologous head, tail, DNA replication, homologous recombination, lysis, integration, and regulatory gene modules, which are arranged in the same genomic order as in lambda, supporting their evolutionary relatedness (**Fig S1**). NC-MV is integrated between *E. coli* genes *ybjL* and *rcdA*. NC-Ino is a previously undescribed filamentous *Inoviridae* phage. Some *E. coli* genomes (*e.g.,* strain SF-468, accession No. CP012625) are neatly missing the 9,335 bp Inovirus prophage while other bacteria (*e.g., Yersinia pestis* CO92 and *Salmonella enterica* AR_0127 as well as *Shigella*, *Citrobacter* and *Proteus* strains) carry very similar *Inoviridae* prophage elements in different genomic contexts.

**Table 1.** Prophages in the *E. coli* NC101 chromosome.

| Prophage | NC101 (locus tag) range | Prophage size (bp) | Phage morphotype | Comments |
| --- | --- | --- | --- | --- |
| NC-SV | ECNC101_17984 - _18284 | 46,386 | Siphovirus | Related to dsDNA phage lambda |
| NC-MV | ECNC101_12352 - _12582 | 35,247 | Myovirus | Related to dsDNA phage P2 |
| NC-Ino | ECNC101_16337 - _16297 | 9,335 | Inovirus | Filamentous ssDNA phage |

**Figure S1.**
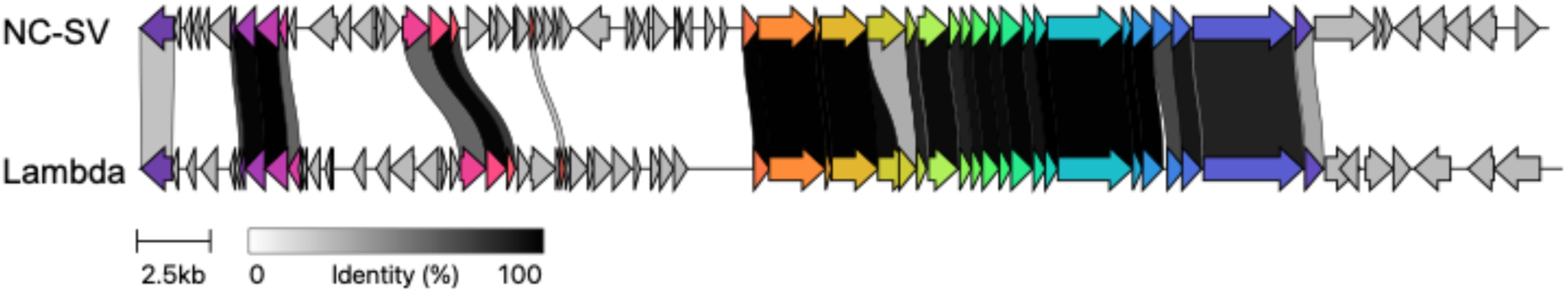
Comparative analysis of NC-SV and lambda prophages. Genomic alignment between the NC-SV prophage and phage Lambda. Arrows represent predicted open reading frames, colored by functional module. Shading between genomes indicates regions of nucleotide identity (0–100%).

Prophages frequently encode mechanisms that protect their bacterial host from infection by competing phages [5, 8]. We hypothesized that one or more of the intact prophages listed in **Table 1** would protect NC101 from lytic phage infection. To test this hypothesis, we deleted each prophage from the NC101 chromosome and challenged with lytic phages isolated from wastewater (phages RSB01–04). Genomic analyses of the RSB wastewater phages reveal that RSB01 and RSB03 are both Veterinaerplatzviruses that share 51.7% nucleotide homology while RSB02 and RSB04 are related Kuraviruses that share 60.7% nucleotide homology (**Fig S2**). Kuraviruses are exemplified by the lytic *E. coli* phage øEco32, whose lifecycle has been characterized [35, 36], and are phylogenetically related to Veterinaerplatzviruses, which form a distinct but allied clade within the same viral order.

**Figure S2.**
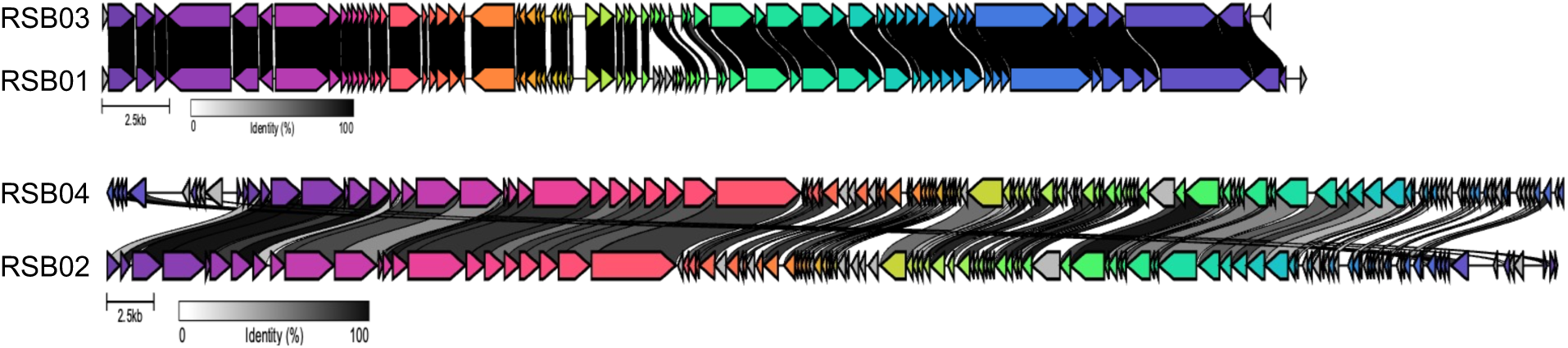
RSB wastewater phage genome comparisons. Phages RSB01-04 were isolated from the headwaters of the wastewater treatment plant in Missoula, Montana, USA. Phage RSB01 (44,523 bp) and RSB03 (43,192 bp) are both Veterinaerplatzviruses that share 51.7% nucleotide homology while phages RSB02 (76,472 bp) and RSB04 (76,787 bp) are related Kuraviruses that share 60.7% nucleotide homology. The genome comparisons shown here were generated by Clinker and show the percent identity of proteins encoded by open reading frames in each phage genome.

Sensitivity to lytic phage infection remained unchanged in NC101^ΔNC-Ino^ and NC101^ΔNC-MV^ (**Fig 1A, Fig S3A**). However, plaque size for lytic phages RSB01 and RSB03 was significantly larger in NC101^ΔNC-SV^ compared to wild-type cells, while no significant differences were observed for RSB02 and RSB04 in any strain tested (**Fig 1A and B, Fig S3A-D**). In liquid culture, RSB01 and RSB03 were also more virulent against NC101^ΔNC-SV^ than against wild-type NC101 (**Fig 1C**), whereas no differences in virulence were observed for RSB02 and RSB04 in any strain tested (**Fig S3E-G**). Additionally, we did not observe any growth defects in uninfected cultures of any prophage mutant compared to wild-type NC101 (**Fig 1C and Fig S3E-G**). These findings suggest that the NC-SV prophage encodes a defense mechanism effective against specific lytic phages.

**Figure 1:**
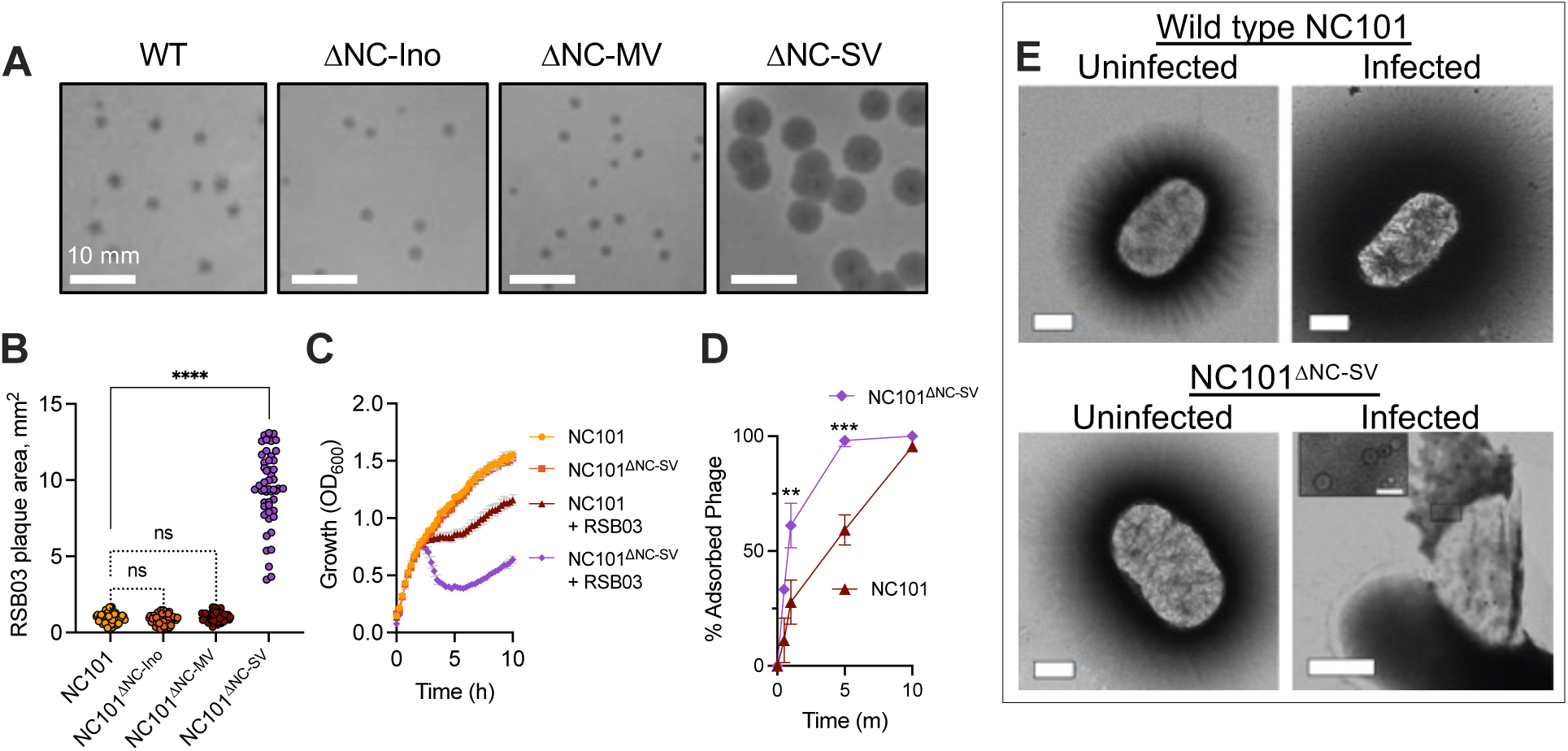
The NC-SV prophage protects *E. coli* NC101 from lytic phage infection by reducing virion adsorption. **(A)** Plaques formed by wastewater phage isolate RSB03 on *E. coli* NC101 or the indicated prophage mutants. Representative images are shown. Scale bars represent 10 mm. **(B)** The surface area of RSB03 plaques formed on lawns of the indicated strains was measured, N=50 plaques per condition, ****P<0.0001 compared to NC101; ns, not significant. **(C)** Growth (OD_600_) of NC101 and NC101^ΔNC-SV^ was measured after infection with RSB03 (MOI=1). **(D)** The percentage of adsorbed RSB03 virions was measured in NC101 or NC101^ΔNC-SV^ cells at the indicated times post infection, MOI=1. Data are the mean ± SEM of three experiments, **P<0.01, ***P<0.001. **(E)** Representative transmission electron microscopy images of negatively stained cells of the indicated strains 60 minutes post-infection with phage RSB03 (MOI 1). Scale bars represent 500 nm (50 nm in the inlay).

An increase in plaque size for phages RSB01 and RSB03 but not RSB02 or RSB04 suggests that a cell surface receptor used by RSB01 and RSB03 may be differentially regulated by NC-SV, which could affect phage adsorption to host cells. To test this, we performed phage adsorption assays on NC101 and NC101^ΔNC-SV^. Phage RSB01 and RSB03 adsorption was significantly (P<0.01) higher in NC101^ΔNC-SV^ compared to NC101 cells (**Fig 1D, Fig S3H**) while adsorption of phages RSB02 and RSB04 was not affected (**Fig S3I and J**), indicating that RSB02 and RSB04 use a different cell surface receptor than RSB01 and RSB03. Examination of RSB03-infected cells by transmission electron microscopy revealed that NC101^ΔNC-SV^ cells appear more prone to lysis by RSB03 compared to NC101 cells (**Fig 1E**). Overall, these data indicate that the NC-SV prophage regulates a cell surface factor to reduce adsorption of some phage types to *E. coli* NC101.

**Figure S3.**
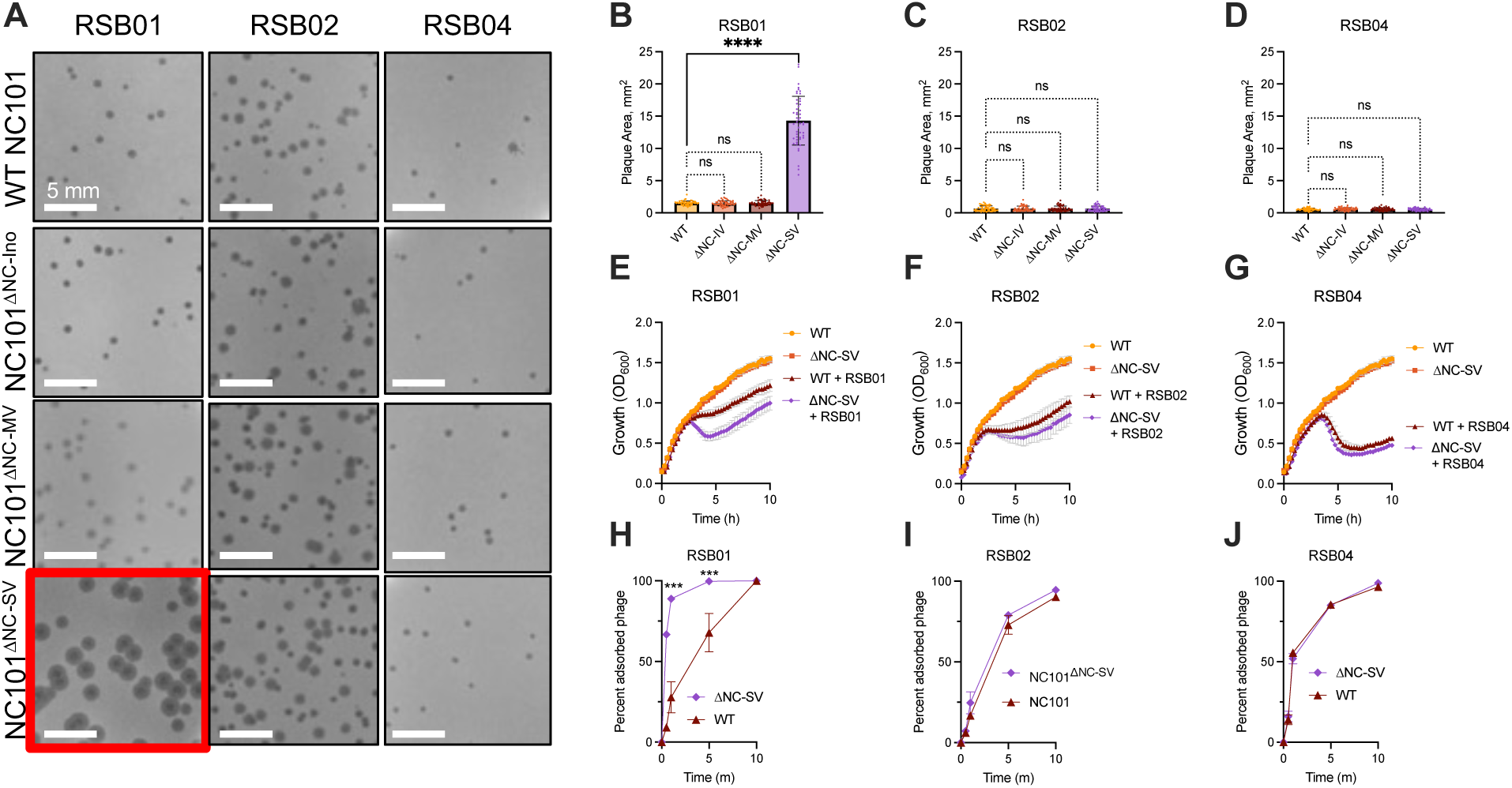
The NC-SV prophage protects *E. coli* NC101 from infection by some, but not all lytic phages by reducing virion adsorption. **(A)** Representative images of plaques formed by wastewater phage isolates RSB01, RSB02, or RSB04 on WT NC101 or the indicated prophage mutants are shown. **(B-D)** The surface area of plaques formed on lawns of the indicated strains was measured, N=50 plaques per condition, ****P<0.0001; ns = not significant. **(E-G)** Growth of WT NC101 and NC101^ΔNC-SV^ was measured after infection with the indicated phages at an MOI of one. **(H-J)** The percentage of adsorbed virions was measured in WT NC101 or NC101^ΔNC-SV^ cells at the indicated times post infection, MOI=1. Data are the mean ± SEM of three experiments, ***P<0.001.

### The NC-SV prophage restricts lytic phage replication in the mouse intestine

Although prophage mediated inhibition of infection by competing phages is well characterized *in vitro*, infection dynamics in more complex ecological systems such as the gut are less well characterized. To test activity of RSB03 in the mouse intestine, we colonized specific pathogen-free C57BL/6 male mice with NC101 and administered RSB03 or heat-inactivated RSB03 enterally (**Fig S4A**). Fecal *E. coli* densities remained stable and comparable between groups (**Fig S4B**), indicating that acute RSB03 administration did not affect bacterial colonization density. RSB03 titers in fecal pellets were significantly higher and more sustained in NC101-colonized mice that received viable RSB03 compared to those given heat-killed RSB03. In mice given viable RSB03, PFUs were detected transiently shortly after administration and again between days 9 and 12 (**Fig S4C**). These results are consistent with lytic replication of RSB03 in the intestinal environment.

**Figure S4.**
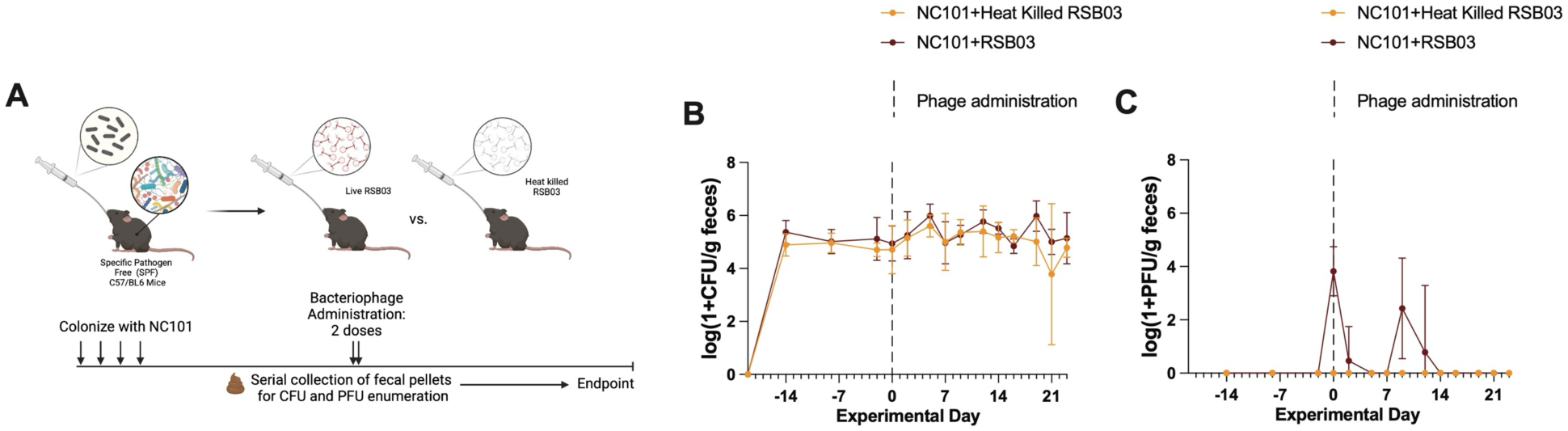
Lytic phage RSB03 replicates in the mouse gut. **(A)** Schematic overview of the experimental design. Specific pathogen-free (SPF) C57BL/6 mice were stably colonized with NC101 *E. coli* followed by oral administration of either active or heat-inactivated RSB03 phage (3×10^7^ PFU per dose, 2 doses, 7-hours apart). **(B)** Qualification of *E. coli* colony forming units (log_10_ CFU/g feces) from fecal pellets. **(C)** Quantification of phage plaque-forming units (log_10_ PFU/g feces) from fecal pellets; mice that received active RSB03 had significantly higher detectable PFU (P<0.0001) which was transiently detected the day of and up to 12 days after administration.

Building on our in vitro findings, we hypothesized that the NC-SV prophage protects *E. coli* NC101 from lytic phage infection *in vivo*. To test this, germ-free C57BL/6 mice were colonized with NC101 or NC101^ΔNC-SV^ and treated daily with RSB03 phage for five days (**Fig 2A**, **Fig S5A**). Fecal *E. coli* density dropped approximately 2.6 log₁₀ CFU/g during treatment but remained stable thereafter in both groups, with no significant differences observed at either day 4 or day 15 (**Fig 2B–C**), consistent with endpoint tissue CFUs in the small intestine shown in **Fig S5B**.

**Figure 2.**
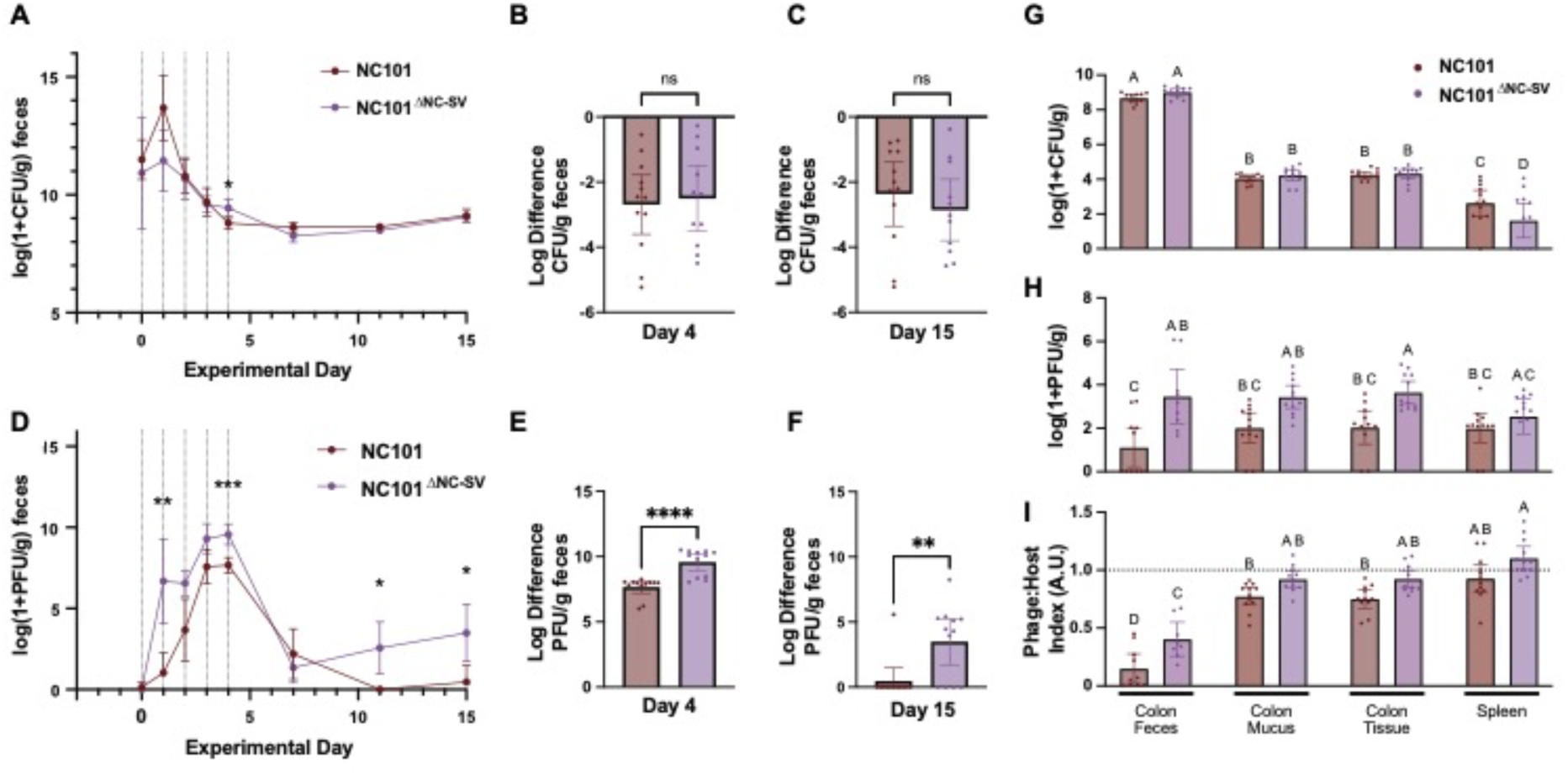
The NC-SV prophage restricts RSB03 phage replication and persistence in the mouse gut. **(A)** Serial quantification of *E. coli* log_10_ CFU from fecal pellets, normalized to fecal sample weight (g). Vertical dotted lines indicate days of RSB03 administration. **(B)** Log_10_ difference in *E. coli* density in feces from baseline to day 4 (last day of RSB03 treatment). **(C)** Log_10_ difference in *E. coli* density in feces from baseline to day 15. (**D**) Serial quantification of plaque forming units (log_10_ PFU) from fecal pellets, normalized to fecal sample weight (g); *p*<0.0001. **(E)** Log_10_ difference in PFU from fecal pellets from baseline to day 4. **(F)** Log_10_ difference in PFU from fecal pellets from baseline to day 4. **(F)** Log_10_ difference in PFU from fecal pellets from baseline to day 15. (**G-**I) Detectable *E. coli* (G) log_10_ CFU, (H) RSB03 PFU, and (I) Phage:Host index min-max normalized log_10_ difference of PFU-CFU, with 1 reflecting equal density for the indicated samples at endpoint (day 17). For bar plots, individual subjects are represented by dots overlaying the mean +/− 95% confidence interval. Significantly different groups are indicated by symbols or compact letter display. ns, not significant, *P<0.05, **P<0.01, ***P<0.001, ****P<0.0001.

In contrast, phage titers in fecal pellets were significantly higher in NC101^ΔNC-SV^ colonized mice during and after treatment, with RSB03 detectable in most NC101^ΔNC-SV^ colonized mice but only transiently or not at all in those colonized with NC101 (**Fig 2D– F**, **Fig S5C**). At endpoint (day 17), NC101^ΔNC-SV^ mice exhibited higher phage loads throughout the gut, including colon contents, mucus, and tissue (**Fig 2H**), with similar trends in the small intestine observed in **Fig S5C**.

Despite similar bacterial loads in tissue compartments (**Fig 2G**, **Fig S5B**), phage:host ratios were significantly elevated in mucus and tissue niches compared to feces (**Fig 2I**, **Fig S5D**), suggesting these sites may facilitate more efficient phage replication or persistence. Notably, NC101^ΔNC-SV^ mice exhibited higher phage burdens across gut and systemic sites (**Fig 2H–I**, **Fig S5C–D**) despite comparable *E. coli* densities (**Fig 2G**, **Fig S5B**), indicating that NC-SV limits lytic phage replication in the gut and may indirectly influence bacterial dissemination to extraintestinal tissues by modulating phage pressure.

**Figure S5.**
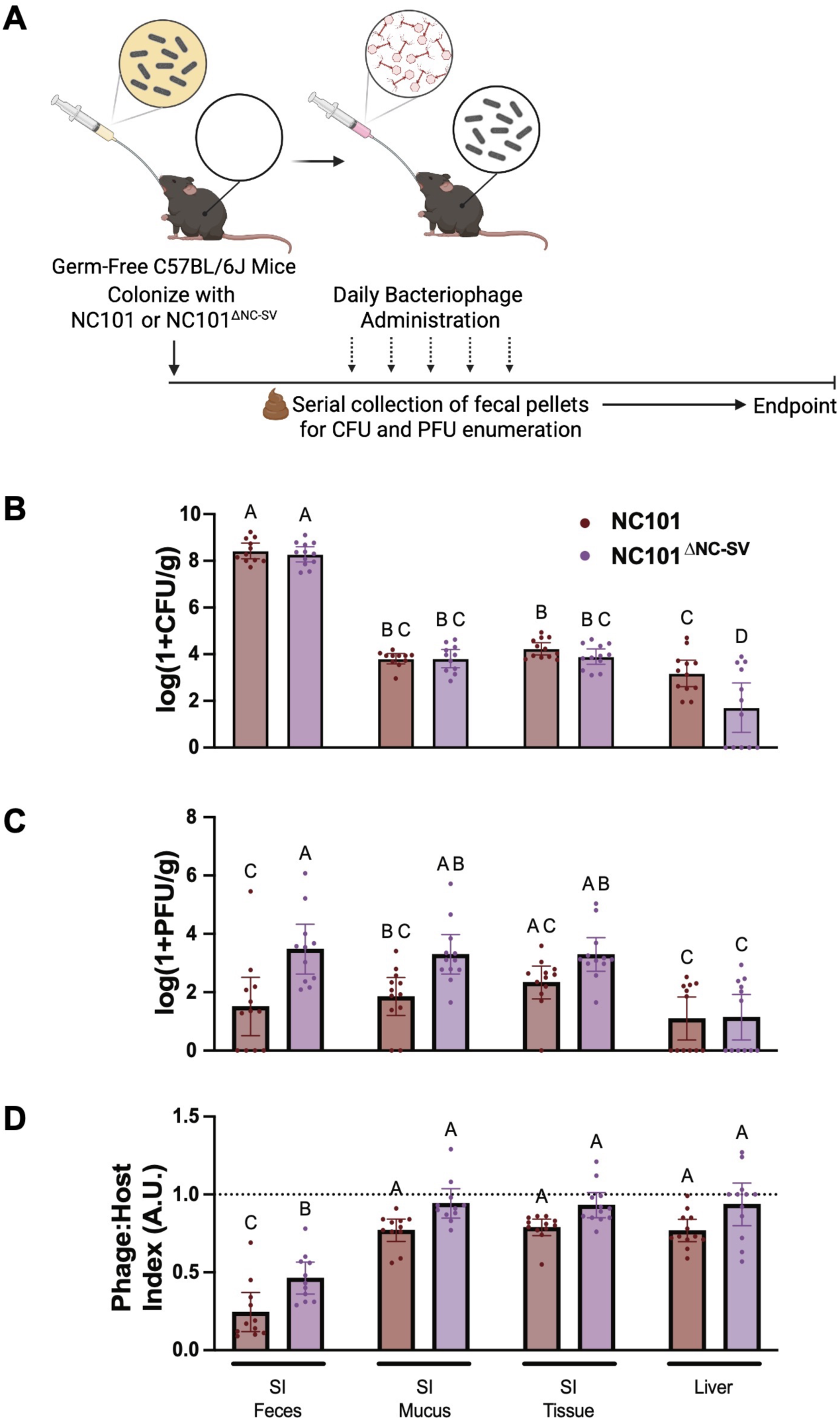
The NC-SV prophage is associated with reduced *E. coli* dissemination to extraintestinal tissues and lower lytic phage loads in the gut. **(A)** Schematic overview of experimental design. Germ-free C57BL/6 mice were stably colonized with either NC101 or NC101^ΔNC-SV^ *E. coli*, then administered 1×10^8^ PFU RSB03 daily by gavage for five consecutive days. Fecal pellets, luminal contents, and tissues were collected for quantification of colony-forming units (CFU) and plaque forming units (PFU). (**B-D**) Detectable *E. coli* CFU (B), RSB03 PFU (C) and Phage:Host index (D, min-max normalized log difference of PFU-CFU, with 1 reflecting equal density) for the indicated samples at endpoint. SI, small intestine. For bar plots, individual subjects are represented by dots overlaying the mean +/− 95% confidence interval. Significantly different groups are indicated by compact letter display.

### The NC-SV prophage downregulates *lamB* and other maltodextrin transport genes and reduces phage adsorption to *E. coli* NC101

To gain a deeper understanding of how the NC-SV prophage affects bacterial responses to lytic phage infection, we performed RNAseq on wild-type NC101 and NC101^ΔNC-SV^ cells ten minutes post-infection with lytic phage RSB03 (MOI=1). Overall, 235 bacterial genes were significantly (P<0.05) differentially regulated at least ±two-fold in NC101 compared to NC101^ΔNC-SV^ (**Fig 3A, Dataset 1**). Gene enrichment analyses revealed that genes associated with carbohydrate transport, amino acid metabolism, and central carbon metabolism were significantly (P <0.05) over-represented (**Fig 3B**).

**Figure 3.**
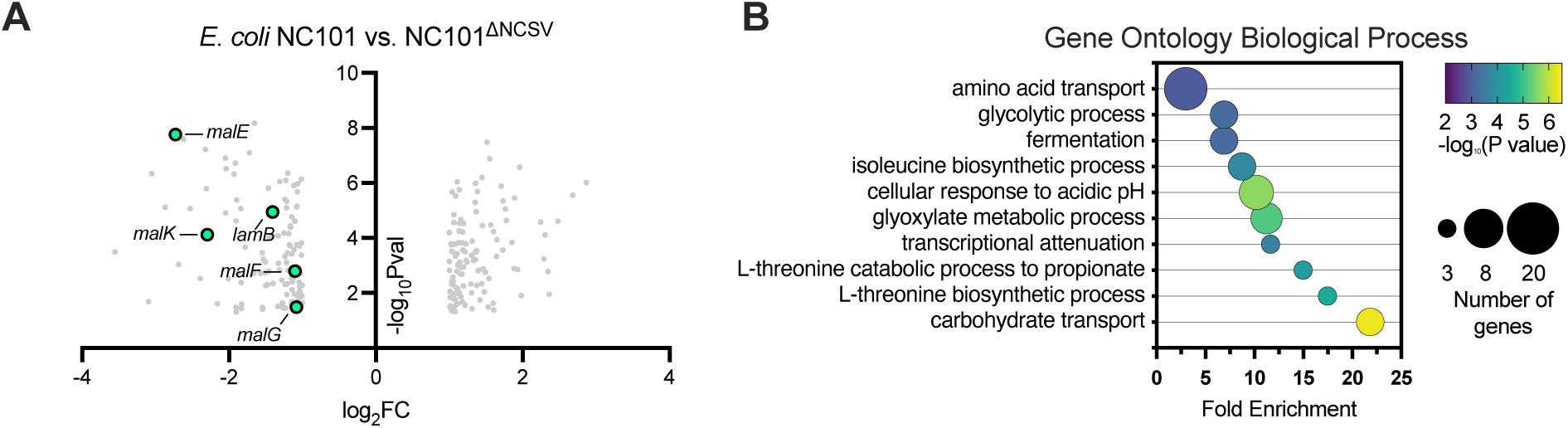
The NC-SV prophage transcriptionally regulates maltodextrin transport and carbon metabolism in response to lytic phage infection. RNAseq was performed on mid-log NC101 or NC101^ΔNC-SV^ cells growing in LB broth 10 minutes post infection with lytic phage RSB03 (MOI=1). **(A)** Volcano plot showing the 235 differentially expressed genes in NC101 compared to NC101^ΔNC-SV^ with a ±2-fold change and P≤0.05. Data are representative of quadruplicate experiments. **(B)** Gene enrichment analysis was performed on significantly differentially regulated genes shown in panel A.

The downregulation of maltodextrin transport genes (*lamB, malE, malF, malG, malK,* **Fig 3A**) by NC-SV hints at a possible mechanism for reduced phage adsorption. The maltodextrin transporter LamB, which is named after <u>lamb</u>da phage, is used as a receptor by numerous coliphages including lambda, K10, TP1, Φ21, and Bp7 [21, 22, 37–39]. Additionally, LamB expression and accessibility on the cell surface directly influences the efficiency of lambda phage adsorption to *E. coli* [40], making it a logical target during phage-bacteria interactions.

We hypothesized that if NC-SV is regulating LamB expression to reduce phage adsorption to *E. coli* NC101, then lambda phage should exhibit infection phenotypes similar to those of phage RSB03 on *E. coli* NC101. However, lambda failed to infect *E. coli* NC101. This outcome might be due to mutations in the NC101 LamB protein that are localized to its extracellular region—the primary site of phage interaction [41] (**Fig S6**).

**Figure S6.**
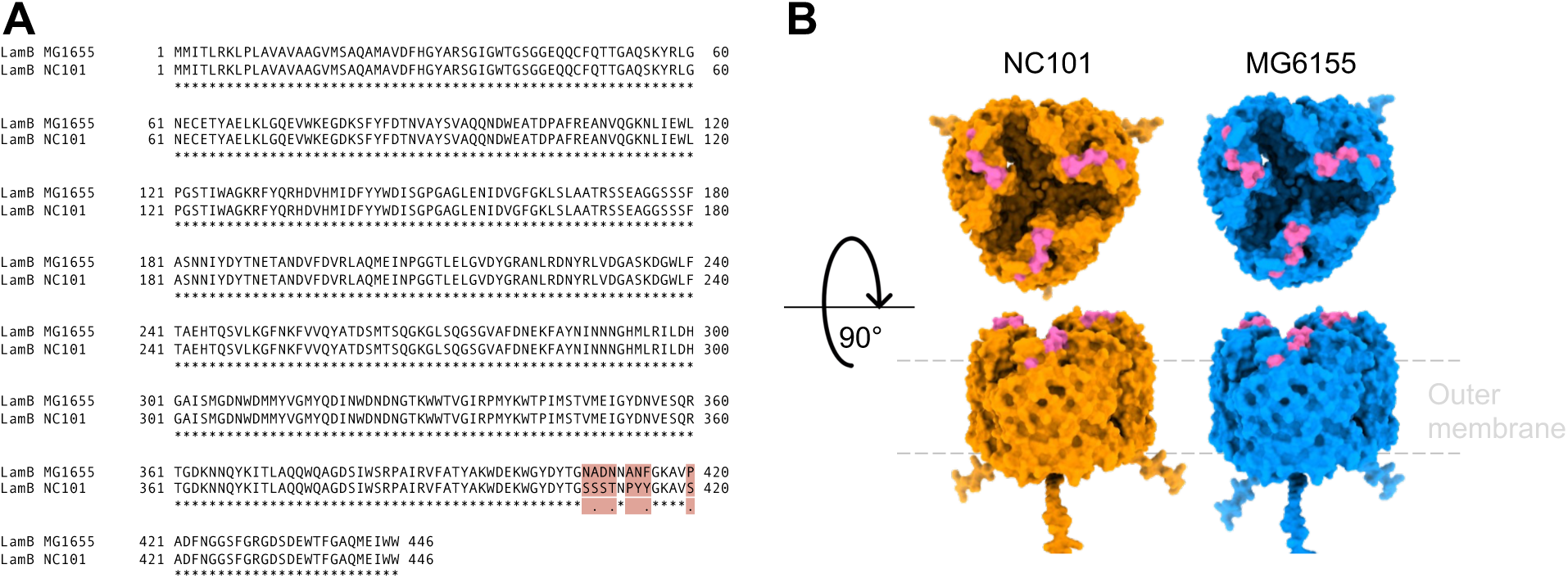
LamB sequence and structure comparison between *E. coli* NC101 and MG6155. **(A)** LamB protein sequences were aligned with Clustal. Differences between the two sequences are highlighted in pink. **(B)** AlphaFold protein predictions of LamB trimers from *E. coli* NC101 and MG6155 are shown in orange and blue, respectively. Residues that are different in NC101 and MG6155 LamB proteins are highlighted in pink.

LamB transports maltodextrin, a glucose polymer, across the outer membrane and into the cell. While LamB is best characterized for transporting maltodextrins— including maltose, maltotriose, and longer linear α-1,4-linked oligosaccharides—it can also facilitate the passage of some other linear oligosaccharides. However, it does not effectively transport monosaccharides like glucose, nor more complex or branched disaccharides and oligosaccharides such as sucrose, lactose, or raffinose, due to structural incompatibility with its narrow, hydrophilic channel [42]. When glucose monomers are abundant, *E. coli* downregulates maltodextrin transport genes via catabolite repression [28]. Consequently, LamB expression decreases, limiting superfluous maltodextrin import and also reducing the number of LamB receptors available for phages [40]. In contrast, when glucose is scarce but maltodextrin polymers are present, LamB surface expression is upregulated [20], increasing phage adsorption to *E. coli* [40].

If NC-SV is regulating LamB expression to reduce competing phage adsorption to *E. coli* NC101, then glucose should reduce RSB03 virion adsorption while maltodextrin should increase RSB03 adsorption. To test this, we supplemented *E. coli* NC101 with 40 mM glucose or maltodextrin. These concentrations were selected because they are comparable to those observed in the intestinal lumen [14, 43, 44]. After 30 minutes, NC101 lawns were infected with lytic phage.

On lawns of NC101 supplemented with glucose, RSB03 plaque size was significantly (P<0.0001) smaller compared to unsupplemented controls (**Fig 4A**). Glucose also reduced RSB03 adsorption to NC101 cells (**Fig 4B**) but had minimal impact on RSB01, RSB02, and RSB04 adsorption (**Fig S7**). Conversely, maltodextrin significantly (P<0.0001) increased RSB03 plaque size on lawns of NC101 (**Fig 4A**), which was associated with an increase in RSB01 and RSB03 adsorption to NC101 cells, but not RSB02 or RSB04 (**Fig 4C; Fig S7**). Finally, sucrose (40 mM), which does not modulate LamB expression [45], had no effect on RSB03 plaque size (**Fig 4A**) nor adsorption to *E. coli* NC101 (**Fig 4D**). Collectively, these results are consistent with the idea that phages RSB01 and RSB03 utilize the maltoporin LamB as a cell surface receptor and that the NC-SV prophage regulates *lamB* expression to defend against competing phages.

**Figure 4:**
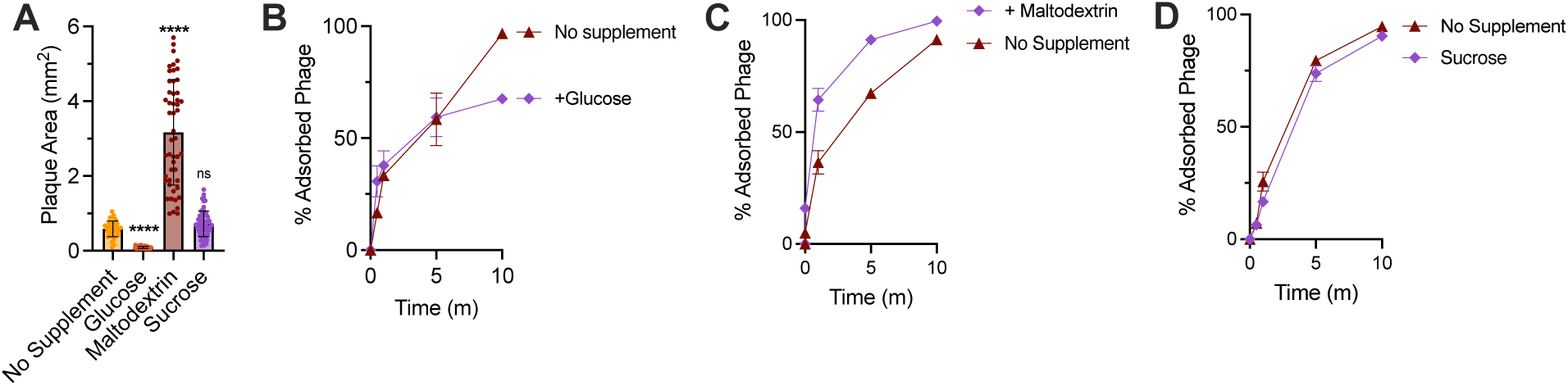
Exogenous sugars modulate phage adsorption to *E. coli* NC101. *E. coli* NC101 was grown in diluted (10%vol/vol) LB broth or agar supplemented with 40 mM of the indicated carbon sources at 37°C. **(A)** RSB03 plaque areas on NC101 lawns were measured after 18 hours. N = at least 30 plaques per condition; ****P<0.0001 compared to the no supplement control; ns, not significant. **(B-D)** The percentage of adsorbed RSB03 virions were measured in cultures supplemented with 40 mM of the indicated sugar at the indicated times post infection with an initial MOI of one. Data are the mean ± SEM of three experiments.

**Figure S7.**
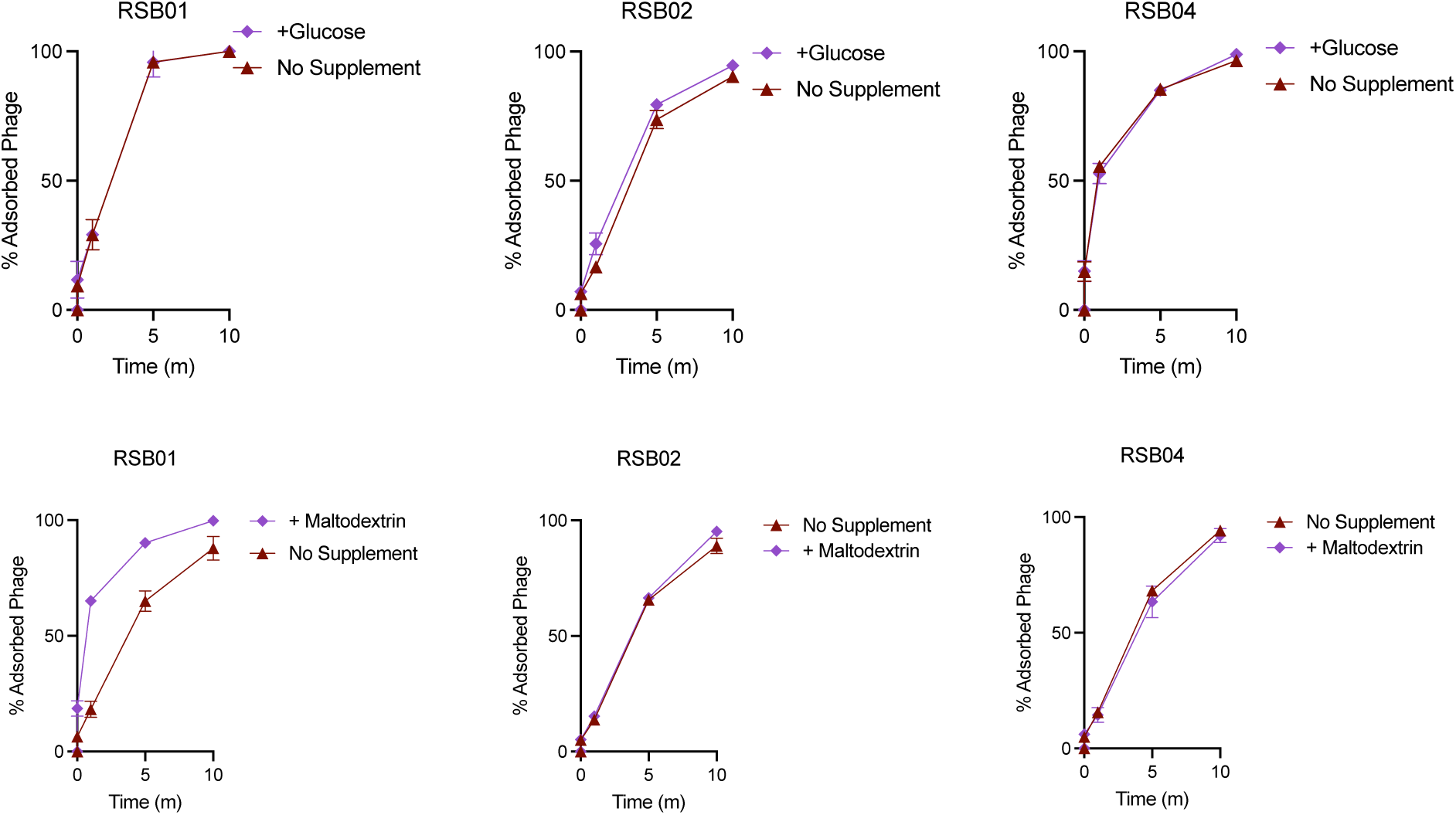
The impact of glucose and maltodextrin on phage virion adsorption to *E. coli* NC101. The percentage of adsorbed RSB03 virions were measured at the indicated times post infection with an initial MOI of one. Data are the mean ± SEM of three experiments.

### An NC-SV-encoded small RNA transcriptionally regulates maltodextrin transport genes and reduces phage adsorption to *E. coli* NC101

Because NC-SV deletion altered competing phage adsorption, we hypothesized that NC-SV encodes gene(s) that influence phage adsorption to *E. coli* NC101. It has been previously shown that another lambdoid prophage, e14, that integrates into the same genomic locus as NC-SV (the *icd* gene) encodes a small RNA (sRNA) called co293 that alters cellular metabolism [46–51]. Specifically, co293 post-transcriptionally regulates the transcription factors HcaR and FadR [46], resulting in the upregulation of propionate degradation pathways and downregulation of glycolysis and the TCA cycle [46, 52].

We hypothesized that NC-SV may encode a co293-like sRNA that modulates bacterial sugar transport and phage adsorption. To test this, we aligned the co293 sRNA sequence from *E. coli* MG1655 to the NC-SV genome and identified a 77-nucleotide region with 58% sequence identity (45/77) located at the 3′ end of the *cI* phage repressor gene on the positive strand (**Fig 5A and B**). The *cI* repressor is a conserved transcriptional regulator in lambdoid phages that maintains lysogeny by repressing lytic gene expression [53]. We refer to this candidate regulatory RNA as *svsR* (NC-<u>SV</u> <u>sR</u>NA). Secondary structure prediction of *svsR* revealed a stable hairpin structure (**Fig 5C**), a common feature of bacterial sRNAs that facilitates RNA stability and target binding [54, 55].

**Figure 5.**
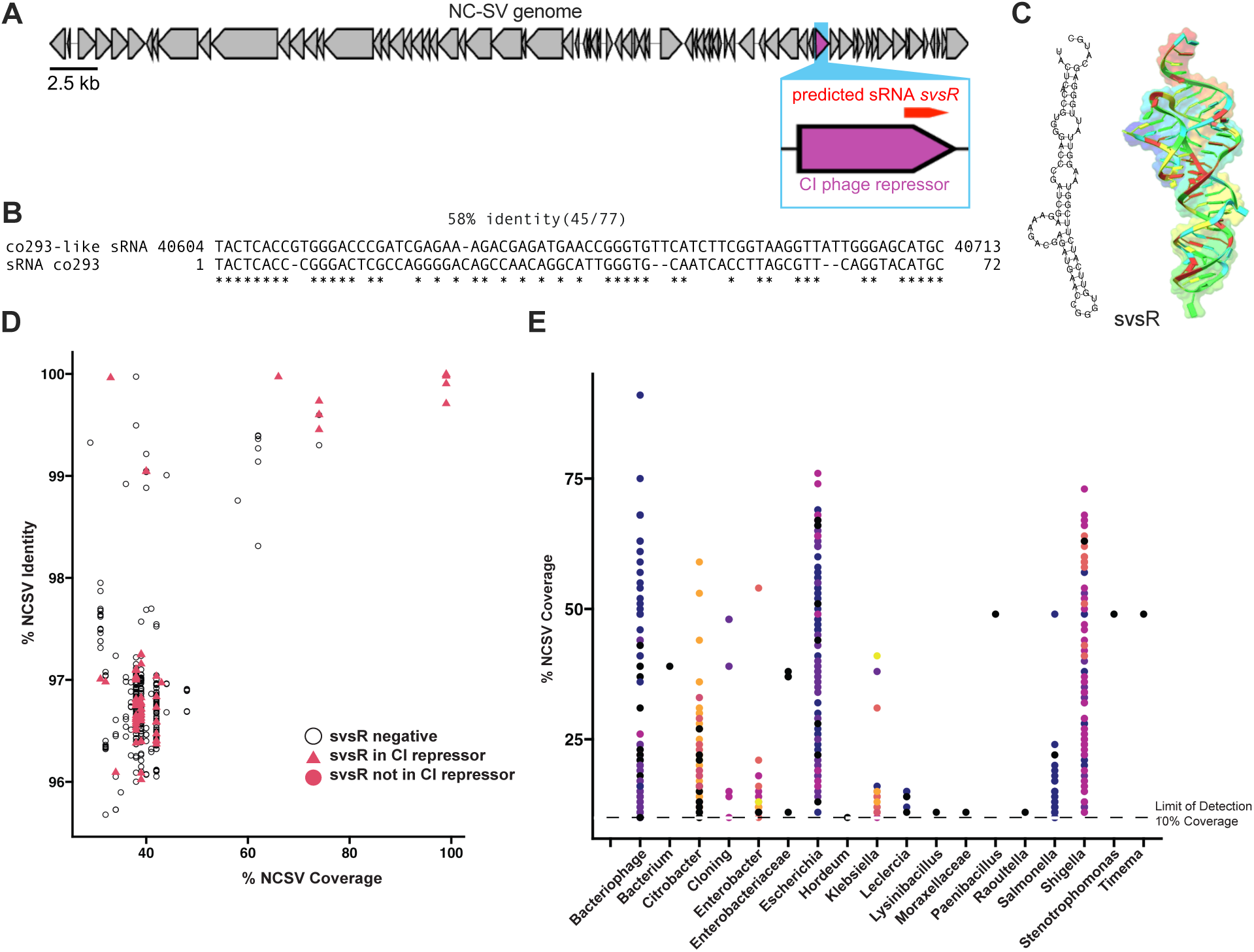
The NC-SV prophage encodes a conserved co293-like small RNA, svsR, found across Enterobacteriaceae. **(A)** Genome organization of the NC-SV prophage. The predicted *svsR* sRNA is encoded on the positive strand at the 3′ end of the *cI* phage repressor gene (highlighted in purple). **(B)** Sequence alignment between *svsR* and the *co293* sRNA from *E. coli* MG1655 reveals 58% identity (45/77 nucleotides). **(C)** Predicted secondary structure (left) and AlphaFold3 structural model (right) of *svsR*, reveal a stable hairpin conformation. **(D)** Conservation of *svsR* among NC-SV–like prophages in *E. coli* genomes with ≥30% NC-SV genome coverage. Each point represents a prophage, with % coverage plotted against % identity to the NC-SV reference. Open circles indicate the absence of *svsR*, while pink symbols indicate its presence; triangle shapes denote *svsR* embedded within the *cI* gene. **(E)** NC-SV– related prophages found in non-*E. coli* Enterobacteriaceae genomes with ≥10% genome coverage. Each dot represents a genome, colored by species or strain.

To evaluate the conservation of *svsR* among related phages, we examined *E. coli* genomes containing NC-SV–like prophages with ≥30% genome coverage. *svsR* was frequently present and often located within the *cI* repressor gene, as indicated by triangle symbols in **Fig 5D**. To determine whether NC-SV-related prophages are also prevalent beyond *E. coli*, we extended our analysis to a broader dataset of publicly available Enterobacteriaceae genomes. We identified NC-SV-like prophage sequences (≥10% genome coverage) in multiple genera, including Klebsiella, *Enterobacter*, *Citrobacter*, and *Salmonella* (**Fig 5E**). These findings suggest that NC-SV-like prophages—and potentially *svsR*—are widespread across the Enterobacteriaceae family.

We next challenged NC101^ΔNC-SV^ expressing *svsR* with lytic phage. Expression of *svsR* did not significantly affect the growth of NC101^ΔNC-SV^ (**Fig 6A**) but did reduce bacterial lysis by RSB03 (**Fig 6B**) and decreased RSB03 plaque size by approximately half compared to empty vector controls (**Fig 6C and D**). Further, the *svsR* sRNA reduced RSB03 virion adsorption to NC101^ΔNC-SV^ cells (**Fig 6E**). These observations demonstrate that *svsR* expression recapitulates the protective effects of the NC-SV prophage.

**Figure 6.**
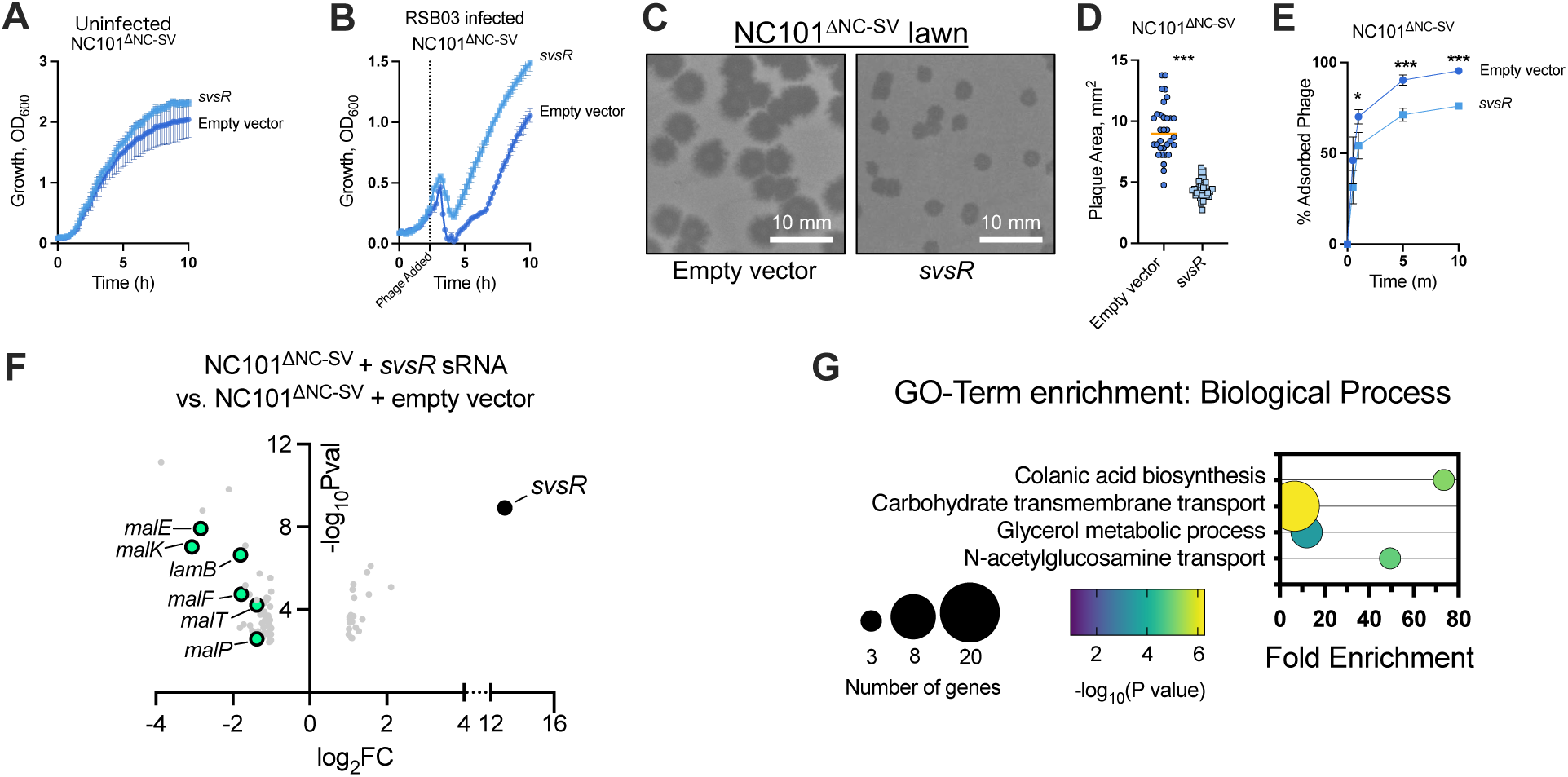
The *svsR* sRNA downregulates maltodextrin transport genes and protects *E. coli* from lytic phage infection. **(A and B)** Growth (OD_600_) of NC101^ΔNC-SV^ carrying an empty expression construct or a *svsR* sRNA expression construct was measured at the indicated times post infection with lytic phage RSB03 at an MOI of one. Data are the mean ± SEM of three experiments. **(C)** Representative images of RSB03 plaques on the indicated lawns. **(D)** RSB03 plaque area was measured after 18 hours on lawns of NC101^ΔNC-SV^ expressing *svsR* in *trans* or empty vector control. **(E)** The percentage of RSB03 virions adsorbed to NC101^ΔNC-SV^ cells carrying the empty vector or the *svsR* expression vector was measured at the indicated times post infection with an initial MOI of one. Data are the mean ± SEM of three experiments; *P<0.05,***P<0.001. (**F**) Volcano plot showing the 83 differentially expressed genes in NC101^ΔNC-SV^ expressing *svsR* compared to the empty vector control with ±2-fold change and P≤0.05. Data are representative of quadruplicate experiments. (**G**) Gene enrichment analysis was performed on the significantly differentially regulated genes shown in panel F.

To determine how *svsR* may affect the bacterial host independent of the NC-SV prophage, we performed RNAseq on NC101^ΔNC-SV^ cells carrying an empty expression vector or the *svsR* sRNA expression vector 10 minutes post infection by phage RSB03. In cells expressing *svsR*, 83 genes were significantly (P<0.05) differentially regulated at least two-fold compared to the empty vector control (**Fig 6F, Dataset 2**). Gene enrichment analyses revealed that genes associated with maltodextrin transport, glycerol metabolism, N-acetylglucosamine transport, and colonic acid biosynthesis (related to biofilm formation in *E. coli* [56]) were significantly (P <0.05) over-represented (**Fig 6G**). Specifically, maltodextrin transport genes (*lamB, malE, malF, malK, malP, malT*) were downregulated, similar to the pattern of maltodextrin transport genes downregulated by the NC-SV prophage (**Fig 3**), most notably the phage receptor *lamB* (**Fig 6F**). Additionally, other carbohydrate transport genes (*fruB, treB, treC, mglC, mglB, manX, manZ, manY, psuT, psuG*) were also downregulated (**Dataset 2**) consistent with broader effects on bacterial host metabolism.

Collectively, these results suggest that NC-SV encodes a co293-like sRNA called *svsR* that downregulates polysaccharide transport genes, including *lamB*, to reduce phage adsorption to *E. coli* NC101.

## Discussion

Our study demonstrates that the lambda-like prophage NC-SV protects adherent-invasive *E. coli* from lytic phage infection by downregulating maltodextrin transport genes, including the phage receptor *lamB*. This regulation reduces virion adsorption to the bacterial surface and limits phage replication both *in vitro* and *in vivo*. These findings provide mechanistic insight into how prophages can modulate bacterial physiology to enhance survival under phage predation, which has implications for AIEC persistence in environments such as the inflamed IBD gut. Notably, NC-SV is also present in the prototypical AIEC strain LF82 [57], suggesting that its phage defense functions may be conserved among clinically relevant AIEC lineages.

While prophage-conferred resistance to lytic phages is a well-established outcome of lysogenic conversion [5], the diversity of mechanisms involved remains underexplored. Our work adds to this body of research by showing that NC-SV confers indirect resistance via transcriptional regulation of host carbohydrate transport systems. By repressing maltodextrin transport genes, NC-SV limits the number of available receptors for phages that rely on LamB for adsorption.

Beyond phage resistance, NC-SV–mediated repression of maltodextrin transport may also modulate AIEC metabolic fitness. Dietary maltodextrins—prevalent in Western diets—enhance AIEC colonization [16] but also induce LamB expression, thereby increasing phage susceptibility [19, 20]. Thus, NC-SV may help AIEC navigate a trade-off between nutrient acquisition and phage vulnerability. This is consistent with emerging evidence linking bacterial metabolism to phage-host dynamics [58–61].

Our *in vitro* findings led us to evaluate whether NC-SV also confers protection in more complex, host-associated environments. Importantly, intestinal *E. coli* loads remained stable and comparable between mice colonized with NC101 and those colonized with NC101^ΔNC-SV^, indicating that NC-SV does not affect bacterial abundance within the intestinal lumen. However, phage titers were significantly higher in feces and intestinal tissues of NC101^ΔNC-SV^ colonized mice, demonstrating that NC-SV restricts lytic phage replication *in vivo*. Notably, elevated phage loads in NC101-colonized mouse intestines correlated with reduced *E. coli* burden in extraintestinal tissues such as the liver and spleen, suggesting that NC-SV-mediated control of phage replication may also enhance bacterial dissemination beyond the gut or persistence in extraintestinal sites in the presence of lytic phage predation.

To identify molecular mechanisms underlying these effects, we examined NC-SV-regulated transcriptional responses during phage infection. Our RNA sequencing data revealed that an sRNA called *svsR* encoded by NC-SV downregulates maltodextrin transport genes, including *lamB*. While the specific mechanism of action of this sRNA remains to be determined, sRNA-mediated gene regulation has emerged as an important mechanism for bacterial adaptation to environmental stressors. For example, in *E. coli* and *Salmonella enterica,* envelope stress induces the σ^E^-dependent pathway, which regulates, among other genes, the expression of sRNAs that cause an abrupt downregulation in the expression of numerous outer membrane proteins [62]. In *S. enterica*, one of these outer membrane proteins is LamB, which is downregulated at the mRNA level by the sRNA *micA* in an Hfq-dependent manner [63].

The *svsR* sRNA is encoded on the 3’ end of the *cI* gene. Notably, bacterial sRNAs are often encoded within diverse genomic contexts, including the 3′ ends of open reading frames. Some 3’ sRNAs are transcribed independently from internal promoters, while others are processed from the 3′ regions of mRNAs by RNase activity. For instance, the s*roC* sRNA in *E. coli* is derived from the 3′ region of the *ygdR* mRNA and plays a regulatory role by acting as an RNA sponge for GcvB [64]. Similarly, *ryhB-2*, a homolog of the well-known iron-responsive sRNA *ryhB*, is transcribed from the 3′ end of the *yhfA* coding region [65]. How *svsR* is transcriptionally regulated or post-transcriptionally processed is presently unclear, and further studies will be necessary to elucidate its precise biogenesis and functional mechanisms. Nonetheless, our findings demonstrate that *svsR* plays a key role in downregulating a common phage receptor and conferring resistance to lytic phage infection, highlighting its functional importance.

In addition to repressing *lamB* and other maltodextrin transport genes, svsR expression also altered transcription of genes involved in colanic acid biosynthesis (Fig 6G), a pathway known to impact bacterial survival in stressful environments. Colanic acid forms an exopolysaccharide capsule that can physically shield *E. coli* from lytic phage predation by blocking receptor access [66]. Its production is also sensitive to carbon source availability, with glucose suppressing and other conditions enhancing its expression [67]. Moreover, colanic acid has been implicated in increased serum resistance and enhanced survival in host-associated environments [68]. Thus, svsR-mediated modulation of colanic acid biosynthesis may further improve *E. coli* resistance to phage infection and promote survival or dissemination beyond the gut, as observed in our *in vivo* experiments.

While its role in phage defense is clear, the specific targets of svsR remain unknown. For example, the sRNA encoded by the related e14 prophage targets the transcription factors *hcaR* and *fadR*, yet these genes were not differentially expressed in any of our RNAseq datasets, suggesting they are unlikely to be direct targets of svsR. Future studies will be needed to identify the regulatory targets and molecular interactions of svsR, as well as to determine whether it influences additional aspects of bacterial physiology beyond phage defense.

Overall, this work highlights how prophages can rewire bacterial metabolism to influence interactions with phages. Beyond classical phage defense systems (e.g., restriction modification, CRISPR-Cas, etc.), bacteria may exploit prophage-encoded regulatory elements to balance nutrient acquisition with phage resistance. These strategies likely support bacterial persistence in dynamic environments like the gut, where microbial communities are continually shaped by diet, interaction with the mammalian immune system, and phage predation. A deeper understanding of prophage-driven metabolic regulation could ultimately guide the development of dietary or pharmacological interventions to modulate microbial ecosystems and mitigate dysbiosis.

## Methods

### Bacterial and Phage Strains, Growth Conditions, Plasmids, and Primers

Bacterial strains, phages, plasmids, and their sources are listed in **Table 2**. Primer sequences are listed in **Table 3**. Unless indicated otherwise, bacteria were grown in LB at 37°C with shaking and supplemented with antibiotics or 0.1% arabinose when appropriate. Unless otherwise noted, antibiotics were used at the following concentrations: gentamicin (10 or 30 µg mL^−1^), ampicillin (100 µg mL^−1^), and carbenicillin (300 µg mL^−1^).

**Table 2.** Bacterial strains, phages, and plasmids.

| Strain | Source |
| --- | --- |
| NC101 | [24] |
| NC101<sup>ΔNC-Ino</sup> | This study |
| NC101<sup>ΔNC-MV</sup> | This study |
| NC101<sup>ΔNC-SV</sup> | This study |
| **Phages** |  |
| RSB01 | This study |
| RSB02 | This study |
| RSB03 | This study |
| RSB04 | This study |
| **Plasmids** |  |
| pSLTS | Addgene, [69] |
| pT2SK | Addgene, [69] |
| pHERD20T | [70] |
| pHERD20T-sRNA | This study |

**Table 3.**
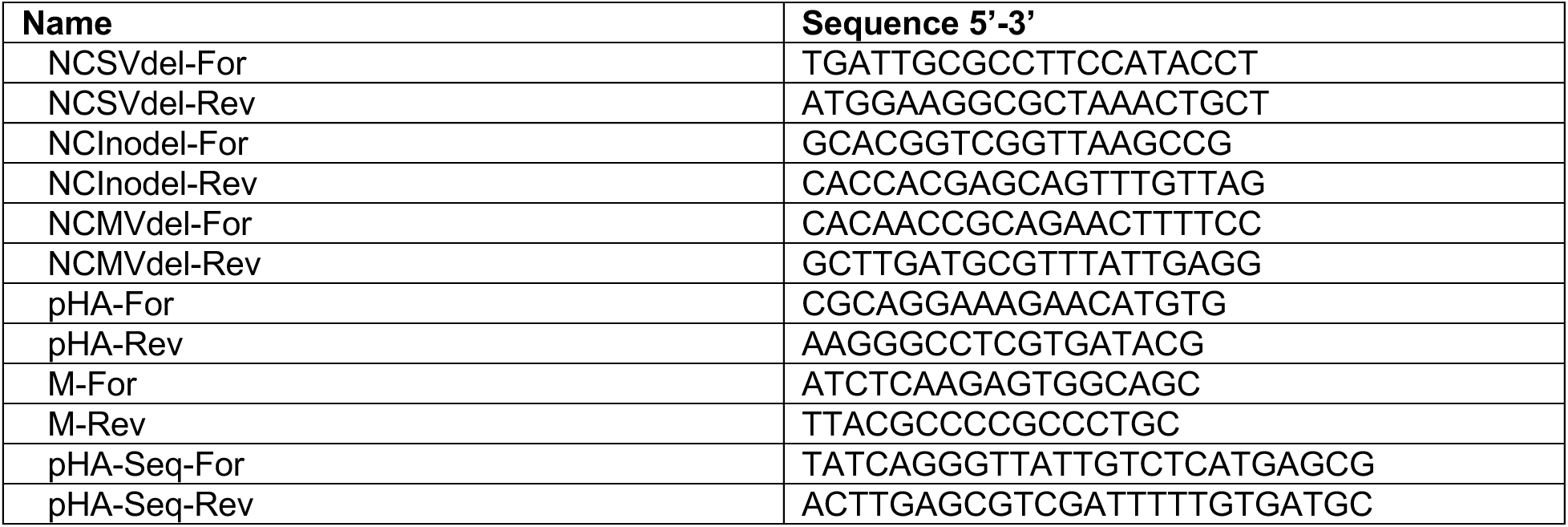
Primers.

### Wastewater phage isolation and sequencing

Samples were collected from the headwaters of the Missoula, Montana wastewater treatment plant. Collected samples were then plated onto lawns of *E. coli* NC101. Individual plaques were picked, plaque purified, and propagated in NC101 growing in LB broth. DNA was isolated from plaque purified phage isolates and sequenced by SeqCoast Genomics, LLC (Portsmouth, NH, USA). Phage genomes were assembled by SPAdes [71] and visualized using clinker [72].

### Construction of NC101 prophage mutants

Prophages NC-Ino, NC-MV, and NC-SV were individually deleted from the NC101 genome using pSLTS mediated genome editing [69]. Briefly, 5’ and 3’ ends of prophage mutation cassettes were designed with homology regions upstream and downstream of each prophage to be deleted and ordered as gene blocks (Azenta). Completed mutation cassettes for each prophage were constructed through Gibson Assembly using the primers listed in Table 3 to fuse 5’ and 3’ ends to a I-SceI cleavable kanamycin selectable marker and plasmid backbone from plasmid pT2SK. Completed prophage mutation cassettes were then introduced to NC101 cells containing the pSLTS helper plasmid via electroporation. pSLTS containing NC101 cells were induced with 2 mM L-arabinose to induce the expression of the lambda red recombinase to aid in recombination of mutation cassettes into the NC101 genome. Resulting colonies were screened for kanamycin resistance and sequenced to confirm successful mutation cassette integration. Once prophage deletions were confirmed, the kanamycin resistance marker was removed by plating PBS-resuspended colonies onto LB-amp plates containing 100 µg/mL anhydrotetracycline to induce the expression of the pSLTS encoded I-SceI endonuclease. Resulting colonies were then sequenced to confirm scarless deletions of each NC101 prophage.

### Plaque assays and plaque surface area measurements

Plaque assays were performed using lawns of the indicated strains grown on the indicated plates. Phages in filtered supernatants were serially diluted 10× in PBS and spotted onto lawns of the indicated strain. Plaques were imaged after 18 h of growth at 37°C. Plaque surface area measurements were performed on individual plaques with image J.

### Growth curves

Overnight cultures of the indicated strains were diluted to an OD_600_ of 0.05 in 96-well plates containing LB and, if necessary, the appropriate antibiotics. After 3 h of growth at 37°C, strains were infected with the indicated phage and growth measurements resumed. OD_600_ was measured using a CLARIOstar (BMG Labtech) plate reader at 37°C with shaking prior to each measurement.

### Phage adsorption assays

Phage adsorption assays were performed as previously described [73]. Briefly, A 18 mL test tube containing 5 mL of LB was inoculated with 100 µL of an overnight culture of the indicated strain. The culture was incubated at 37°C with shaking at 200 RPM for 1 hour until it reached a density of approximately 1 × 10^8^ cells/mL (OD ∼ 0.2). Fresh lysate containing 1 × 10^8^ phages (N_total_) was then added, and the mixture was incubated at 37°C without shaking. At the indicated times, 500 µL aliquots were collected and centrifuged at 4,000g for 1 minute at room temperature to pellet adsorbed phages. Dilutions of the supernatant from the centrifuged sample (representing unabsorbed phage) were plated to determine phage titers. The total phage count (N_total_) and the free phage count (N_free_) were calculated from plate counts using the equation: N_adsorbed_ = N_total_ – N_free_.

### Nutrient supplementation assays

LB broth was diluted with water to 10% vol/vol. For experiments where LB agar plates were used, the dilute LB was supplemented with agar (1.5% w/vol). Glucose or maltodextrin was then supplemented to the dilute LB broth or LB agar to a final concentration of 40 mM. Plaque assays or phage adsorption assays were then performed as described above.

### Mouse model of intestinal adherent-invasive *E. coli* infection and enteral RSB03 treatment

NC101 colonization and RSB03 treatment of SPF mice. Two cohorts of 6-week-old male specific pathogen free C57BL/6 mice (HK *n*=5, active RSB03 *n*=5) were ordered from Jackson Labs and housed in conventional cages with 5 mice per cage. 1×10^8^ CFU *E. coli* were administered to each mouse via oral gavage for four consecutive days (days −29 to −26). Fecal pellets were collected once weekly following colonization to assess baseline *E. coli* density and confirm lack of baseline lytic phage activity in fecal pellets. On day 0, 3×10^7^ PFU of RSB03 (or an equivalent volume of the heat-inactivated RSB03 from the same preparation) were administered by oral gavage in two doses, 7 hours apart. Fecal pellets were collected prior to the second dose of RSB03, then every 2-3 days for CFU and PFU quantification.

For *E. coli* monoassociation and RSB03 treatment of germ-free mice. Two cohorts each containing 12 mice (6 female, 6 male) were housed with 2-4 mice per cage. Germ-free C57BL/6 mice were removed from germ-free isolators and placed into sterile HEPA filtered cages at 6-8 weeks of age. On day 3, each mouse was gavaged with 1×10^8^ *E. coli* (either NC101 or NC101^ΔNC-SV^) as a single dose. Fecal pellets were collected three days following *E. coli* inoculation to quantify baseline *E. coli* density in feces and to confirm lack of baseline lytic phage activity in fecal pellets. From day 0 to day 4, RSB03 (1×10^8^ PFU) was administered daily by oral gavage for five total doses. Fecal pellets were collected daily for *E. coli* CFU and PFU quantification prior to daily oral phage gavage. Additional fecal pellets were collected for analysis on experimental days 7, 11, and 15 following completion of RSB03 treatment. Intestinal contents and tissues, liver, and spleen were collected for CFU and PFU quantification on day 17.

### Quantification of CFU and PFU from mouse fecal pellets, mucus, and tissue samples

Fecal pellets were longitudinally collected from live mice. Intestinal luminal contents from the colon and small intestine were removed following euthanasia for analysis at endpoints. Fecal samples were homogenized in HBSS with Ca^2+^/Mg^2+^ (Corning 21-023-CV) by pipet disaggregation and vortexing for 30 seconds, centrifuged at 50 x g for 10 minutes to pellet fecal debris, and the supernatant was removed (fecal wash). Fecal wash samples were centrifuged at 2,500 x g for 10 minutes to pellet bacteria. The fecal wash supernatant (bacteriophage fraction) was centrifuged at 7,500 x g, 10 minutes, 4°C to pellet any residual bacteria, and supernatant was serially diluted in HBSS with Ca^2+^/Mg^2+^ for PFU quantification. The remaining bacterial pellet was washed with HBSS, then resuspended in HBSS with Ca^2+^/Mg^2+^(bacterial fraction) and serially diluted for CFU enumeration.

Intestinal tissues were thoroughly rinsed three times in PBS. To digest intestinal mucus, 1000 µl of HBSS without Ca^2+^/Mg^2+^ (Corning 21-022-CV) with 10 mM HEPES and 1.5 mM DTT was added to each sample and incubated with shaking for 15 min at room temperature (mucus fraction). Intestinal tissue fragments were removed from mucus digestion and blotted to remove excess liquid. All tissues were homogenized in bead-beating tubes containing two sterile 6.35 mm stainless steel beads (BioSpec) and 0.3% Triton-X / HBSS with Ca^2+^/Mg^2+^ using the Omni Bead Ruptor Bead Mill Homogenizer for 1 min, 4 m/s at room temperature. For PFU quantification, the mucus fraction and tissue homogenate were spun at 7500 x g, 10 min, 4°C to pellet bacteria and debris and supernatant was serially diluted for PFU quantification (phage fraction).

For CFU quantification: The mucus fraction and tissue homogenates were serially diluted 1:10 in HBSS with Ca^2+^/Mg^2+^ and plated on LB-chloramphenicol (30 µg/mL) plates. For fecal samples, 10 µl of each dilution was plated at four 1:10 serial dilutions, and the dilution resulting in countable single colonies was used to determine CFU/g feces.

For PFU quantification: For all phage fractions, 100 µl of the phage fraction and 100 µl NC101 *E. coli* (OD_600_ 0.5-0.7) was added to 5 mL 0.6% soft LB top agar and poured over LB agar plates. Phage fractions were diluted in HBSS with Ca^2+^/Mg^2+^ as needed to attain isolated plaques for counting. Plates were incubated overnight (12-18 hours) prior to counting phage plaques.

### RNA Purification and RNA-seq

Total RNA was extracted from the indicated strains and conditions 10 minutes post phage infection using TRIzol. The integrity of the total cellular RNA was evaluated using RNA tape of Agilent TapeStation 2200 before library preparation. All RNA samples were of high integrity with a RIN score of 7.0 or more. rRNA was first depleted from 500 ng of each sample using MICROBExpress Kit (AM1905, Fisher) following the manufacture’s instruction. The rRNA-depleted total RNA was subjected to library preparation using NEBNext® Ultra™ II RNA Library Prep Kit (E7700, New England Biolabs) and barcoded with NEBNext Multiplex Oligos for Illumina (E7730, New England Biolabs) following the manufacturer’s instructions. The libraries were pooled with equal amounts of moles, further sequenced using MiSeq Reagent V3 (MS-102-3003, Illumina) for pair-ended, 600-bp reads, and demultiplexed using the build-in bcl2fastq code in Illumina sequence analysis pipeline. Raw sequencing reads have been deposited as part of BioProject PRJNA1225266 in the NCBI SRA database.

### RNA-seq Data Analysis

RNAseq data analysis were performed as previously described [74]. Briefly, RNA-seq reads were aligned to the reference *E. coli* NC101 genome (GenBank: GCA_029542525.1), mapped to genomic features, and counted using Rsubread package v1.28.1 [75]. Count tables produced with Rsubread were normalized and tested for differential expression using edgeR v3.34.1 [76]. Genes with ≥twofold expression change and a *P* value below 0.05 were considered significantly differential. Functional classification and Gene Ontology (GO) enrichment analysis were performed using PANTHER classification system (http://www.pantherdb.org/) [77]. RNA-seq analysis results were plotted with GraphPad Prism version 10.

### Structure prediction

AlphaFold3 [78] was used to predict the structures of LamB trimers from *E. coli* NC101 and MG1655 or the putative sRNA encoded by NC-SV. Structures were visualized with ChimeraX-1.9 [79, 80].

### Transmission electron microscopy imaging

Cells were grown to midlog (OD_600_ 0.4), infected with RSB03 (MOI 1) for one hour, washed with PBS, fixed with 4% formamide, and placed on a grid and negatively stained with uranyl acetate. Cells were imaged on a Tecnai Spirit 120 kV TEM.

### Statistical Analyses

Unless specified otherwise, differences between datasets were evaluated by Student’s t test, using GraphPad Prism version 5.0. Longitudinal statistical comparisons between groups from *in vivo* experiments were performed using a mixed effects model with Geisser-Greenhouse correction, with individual timepoints assessed by Sidak’s multiple comparisons test. Mann-Whitney test was used for comparisons between two groups for *in vivo* experiments. Two-way ANOVA with Tukey’s multiple comparisons test was used for statistical comparisons between multiple groups in in vivo experiments. For all tests, a P value < 0.05 was considered statistically significant.

## Supporting Information

**Dataset 1. Differential gene expression in *E. coli* NC101 vs. NC101^ΔNC-SV^ during early lytic phage infection.** RNAseq was performed on mid-log phase NC101 and NC101^ΔNC-SV^ cells grown in LB broth and collected 10 minutes post infection with lytic phage RSB03 (MOI = 1). Dataset includes normalized gene expression values and differential expression statistics.

**Dataset 2. Transcriptomic impact of *svsR* expression in NC101^ΔNC-SV^ during phage infection.** RNAseq was performed on NC101^ΔNC-SV^ cells carrying either an empty vector or the *svsR* sRNA expression construct, collected 10 minutes post infection with lytic phage RSB03 (MOI = 1). Dataset includes normalized gene expression values and differential expression statistics.

## Acknowledgments

This work was supported by NIH grant R01DK124317. P.R.S. was supported by P20GM103474. D.R.F. was supported by the NSF GRFP (366502). N.L.P was supported by the National Center for Advancing Translational Sciences of the National Institutes of Health under Award Numbers UM1TR004409 and K12TR004413 and the Primary Children’s Hospital Foundation Early Career Development Award. The content is solely the responsibility of the authors and does not necessarily represent the official views of the National Institutes of Health or other funding sources.

